# Dynamical insights on the role of supercoiling on DNA radiosensitivity

**DOI:** 10.1101/2025.09.08.674845

**Authors:** Manuel Micheloni, Raffaello Potestio, Lorenzo Petrolli

## Abstract

Ionizing radiation (IR) is a major source of biological hazard, associated with a broad range of detrimental lesions of the structural and molecular integrity of the DNA molecule—often leading to genomic instabilities and severe cellular outcomes.

The radiosensitivity of DNA is deeply affected by a variety of chemical and biophysical factors, which control the dynamical behavior of the molecule as well as diverse cellular processes and response pathways. Among these factors, the role of supercoiling in modulating DNA radiosensitivity remains controversial, with the existing literature being inconclusive on its effective contribution. Here, we characterize the linearization of a supercoiled DNA minicircle by double-strand breaks (i.e., the rupture of the covalent DNA backbone on both complementary strands of the double helix) *via* classical coarse-grained molecular dynamics simulations, and verify how the initial supercoiling regime of the molecule influences the kinetics of the rupturing process. We observe that the excess torsional stress overall enhances the rupturing likelihood but in one specific scenario—associated with a biologically-significant level of (negative) superhelical density: This effect highlights a strong asymmetry between positive and negative supercoiling regimes and provides critical insights on the role of topology on the radiosensitivity of DNA molecules.

## INTRODUCTION

DNA holds the hereditary information required by prokaryotes and eukaryotes to replicate and carry out their biological activity. However, genomic stability is constantly threatened by a variety of toxic agents—associated with both endogenous metabolic processes and exogenous sources. Particularly, ionizing radiation (IR), and the radiolytical by-products thereof, are accountable for a plethora of chemical and structural modifications of the DNA molecule, whose complexity and spatial distribution is determined by the energy and quality of the IR source—and whose biological and mechanical implications involve broad, albeit intertwined, length and time scales [1, 2]. Among the variety of radiation-induced lesions, DNA double-strand breaks (DSBs)—i.e., the disruption of the DNA backbone over both complementary strands—are reckoned be the most cytotoxic, which, if misrepaired or not repaired at all, might lead to severe biological outcomes such as mutagenesis, chromosomal aberrations, apoptosis, and oncogenic transformations [1].

Quantifying the yields and distribution of DSBs at the cellular level, e.g., through *γ*-H2AX immunofluorescence assays [3], have offered a measure of a cell radiosensitivity and capability to handle complex DNA lesions: Yet, this response depends on a complex cascade of enzymatic processes (broadly referred to as the DNA damage and repair machinery), as well as on the morphological modifications associated with defective DNA motif. As for the latter, experimental techniques such as electrophoresis [4] and AFM imaging [5] have dealt critical insights on the stability of the helical structure of DNA upon irradiation [4], and on the diverse biophysical factors affecting the radiosensitivity of the DNA molecule (e.g., the DNA and saline concentrations [6], and the temperature [4]): Particularly, these factors modulate the degree of compaction, as well as active cellular processes involved with the (de)condensation of chromatin [7, 8], thereby influencing the likelihood of structural DNA lesions. Furthermore, a locally-high DNA concentration was shown to act as a radical scavenging mechanisms, thus lowering the yields of strand breaks by radiolytical species, regardless of the irradiation regime [6, 9]: This has been similarly observed by folding DNA into origami nanostructures, despite these frameworks showing a higher sensitivity to dense IR sources [10].

In cells, DNA supercoiling—i.e., the degree of over/underwinding of the double helix about its axis—sets an additional (topological) constraint, known to affect the structural compartmentalization of the molecule as well as key biological processes. It is quantified by the amount of superhelical density (*σ*) [11]:

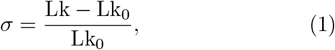

where the linking number (Lk) denotes the number of crossings between the (oriented) strands of a closed DNA molecule—with Lk_0_ defining its reference value (i.e., associated with a torsionally-relaxed molecule). Specifically, negative supercoiling promotes the compaction and local denaturation of DNA in prokaryotes [12], thereby providing a mechanism for gene expression and regulation [13– 15]. Moreover, supercoiling emerges in eukaryotic DNA from its packing about histone cores in nucleosomes, with topoisomerases relaxing the excess torsional stress within key biological processes such as DNA transcription [16]. In fact, the degree of supercoiling modulates the local accessibility of a DNA molecule by shaping its conformational ensemble, potentially exposing specific DNA sequences to exogenous agents [17, 18].

Yet, in spite of its significance, the role of supercoiling on the DNA radiosensitivity is somewhat controversial [19]. Ozols and co-workers [20] observed that the supercoiling hardly influences the yields of DNA strand breaks in regimes of sparse IR, unlike, e.g., the dose-rate and the (concentration of the) buffer, despite their data being admittedly inconclusive. Conversely, Swenberg and Speicher [21] inferred a linear proportionality between the susceptibility and the effective radius of negatively-supercoiled, circular DNA molecules upon irradiation by ^60^Co-*γ* and fission neutron sources—in line with earlier experimental assessments [22, 23]: In fact, the authors demonstrated that the degree of DNA compaction by supercoiling inhibits the induction of (single) strand breaks, an outcome further supported by the standard single-hit target theory. Additionally, the (local and global) structural changes induced by the relaxation dynamics of supercoiled DNA arguably affects the quality, the distribution and the localization of the lesions upon the DNA molecule—according to its co-evolution with the radiation field [24].

In this context, numerical simulations have offered valuable insights on the structural and dynamical evolution of DNA lesions that are hardly accessible by conventional experimental techniques. Monte Carlo track structure (MCTS) codes have been extensively employed to simulate the stochastic sequence of (elastic, inelastic) collisions of the radiation particles within a continuous water-like medium, ultimately depicting the pattern of energy deposition events [25]. Yet, MCTS codes work on static geometric models of the biological targets (e.g., DNA and chromatin fibers), thereby neglecting their inherent dynamics—which is concurrent with the earliest stages of the irradiation process, and whose characteristic timescales are about the same order of magnitude[2]. By characterizing the impact of the IR field [26] and its early-stage effects on DNA [27–29], molecular dynamics (MD) simulations have described the structural and dynamical response of biological systems to molecular lesions, and the repair machinery thereof [30], across multiple resolution levels. In an earlier work, we have carried out a mechanistic and thermodynamic assessment of the DNA rupturing process (driven by diverse DSB motifs) on a biologically-significant timescale, employing a coarse-grained (CG) force field of the DNA double helix [31, 32]: We have shown that the disruption of a DNA molecule is described by a cooperative, thermally-activated process, following the Poisson statistics [33].

In this work, we employ classical MD simulations to extend the characterization of diverse DSB motifs—namely, at a distance *b*_*d*_ = 0, 1, 2, 3 base-pairs (bps) between strand breaks—and verify the impact of DNA supercoiling in modulating the rupturing kinetics of a 672 bp-DNA minicircle. Firstly, we show that mechanically-strained conformations enhance the rupturing probability with respect to a topologically-relaxed DNA molecule: This effect is significantly pronounced at positive supercoiling, highlighting a strong asymmetry between over- and under-coiled topoisomers in the mechanical response to DNA lesions. In addition, we verify that DSB motifs enforced on the tip of the apical loops exhibit a significantly higher rupturing probability than those within the internal supercoiled region of the minicircle, as sharply bent structures are more prone to mechanical failure.

The impact of DNA supercoiling on the linearization kinetics acts in a multiscale manner, and the enhancement effect associated with the DNA rupturing process decreases over time: In fact, a (double) nick of the DNA backbone breaks the topological invariance of the superhelical density, triggering the structural rearrangements of the DNA minicircle driven by the release of the mechanical stress. Finally, we discuss the implications of this work in the context of DNA radiosensitivity. The simulations of DSB motifs enforced in the supercoiled region of DNA minicircles at superhelical density *σ* = −0.06 (which is a scenario of biologically significance [15]) reveals no enhancement in the rupturing probability with respect to a topologically-relaxed scenario: This result was recapitulated by the analysis of the free energy landscapes associated with the DNA rupturing process. Together with the decrease of the effective volume of the DNA target, this observation suggests that this level of supercoiling might effectively lower the radiosensitivity of DNA in cells.

## MATERIALS AND METHODS

### The oxDNA2 force field

All simulations in this work have been performed employing the sequence-dependent (i.e., hydrogen-bonding, stacking) parametrization of the oxDNA2 CG force field, which has been acknowledged to depict the thermodynamic, mechanical, and structural features of the (canonical B-conformation of the) DNA double helix [32, 34, 35]. Originally designed for applications in nanotechnology, oxDNA2 improves on the earliest release of the force field, featuring the major and minor grooves of B-DNA, and an effective electrostatic screening *via* a Debye-Hückel interaction, which refine the behavior of DNA in implicit solvent and at diverse concentrations of monovalent salt—down to a physiological environment [31]. Additionally, by describing nucleotides as dual interaction sites (i.e., involving a backbone and a nucleobase moiety), oxDNA2 offers a convenient resolution tradeoff to capture the dynamical evolution of DSB lesions, while significantly reducing the computational overhead associated with the atomistic description and enabling the exploration of biologically-significant timescales [32, 36].

### Structural characterization of DNA minicircles

Circular DNA molecules show diverse structural dynamics on account of the interconversion between twist and writhe (according to the Călugăreanu-White-Fuller theorem [37, 38]), as well as to the level of superhelical density. Particularly, topologically-constrained DNA minicircles associated with *σ* ≠ 0 exhibit mostly writhed conformations characterized by two structural domains—namely, apical loops, where the DNA helix is sharply bent, and a supercoiled region, where the DNA helix is highly intertwined (see Figure 1).

**FIG. 1.**
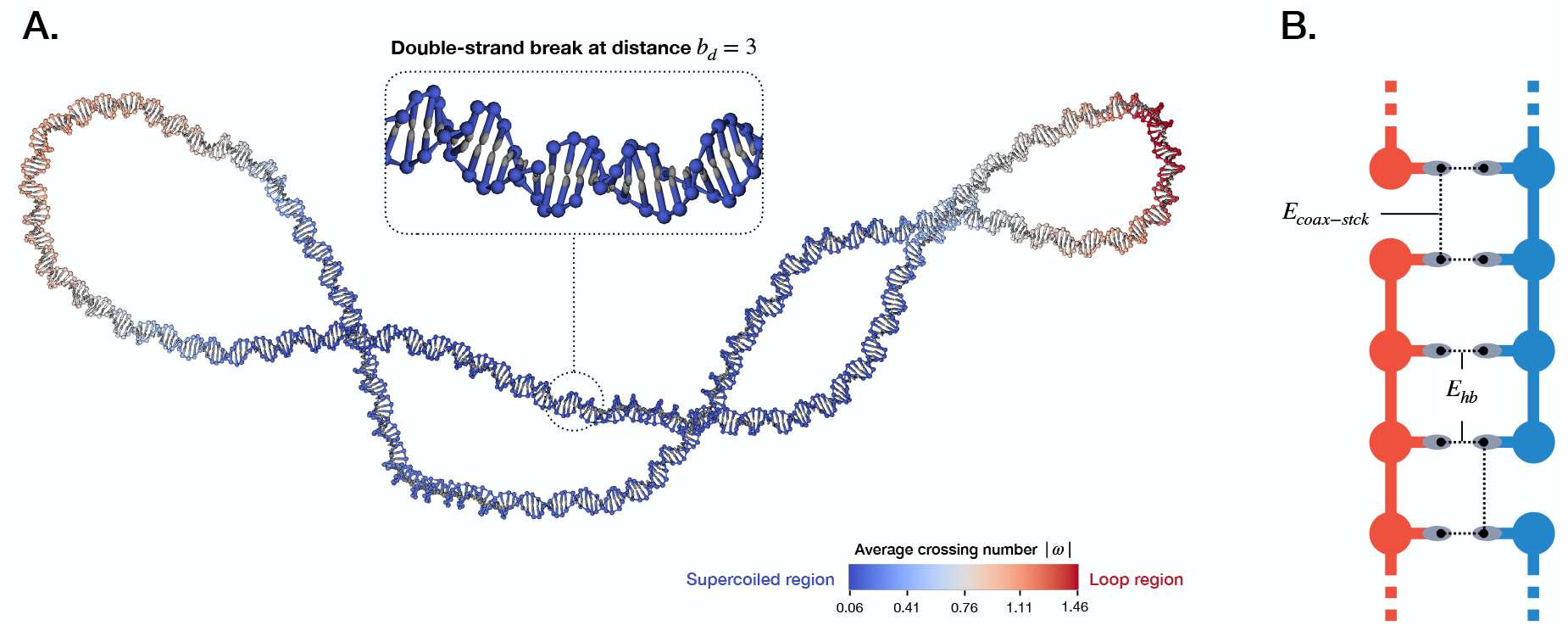
(**A**.) Structure of the 672-bps DNA minicircle employed in this work, broken by a DSB motif of distance *b*_*d*_ = 3. A color scheme has been defined upon the (instantaneous value of the) local writhe associated with each residue. (**B**.) Schematic depiction of a DSB, highlighting the residual (hydrogen-bonding, coaxial-stacking) energy interactions holding the structural integrity of the double helix at the lesion site.

To systematically distinguish between structural motifs of a minicircle along an MD trajectory, we employed the local writhe |*ω*(*i, t*) | [39, 40] (or the average crossing number, ACN [41]), defined as:

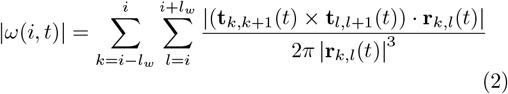

with **r**_*i*_ the coordinates of the *i*-th nucleotide pairing along the centerline of the DNA helix (as such, **r**_*kl*_(*t*) = **r**_*k*_(*t*) − **r**_*l*_(*t*)), **t**_*i,i*+1_ the tangent vector at location (index) *i*, and *l*_*w*_ = 150 bps—corresponding to the persistence length of DNA in solution. From the definition of Equation 2, it follows that high values of the (unsigned) local writhe are associated with highly bent DNA turns, whereas a straight DNA segment contributes zero to the summation.

Thus, the frame-wise calculation of the local writhe lets one monitor the evolution of apical loops and supercoiled regions along the DNA centerline, i.e. by tracking the indices of the nucleotide pairings corresponding to the maxima and minima of |*ω*(*i*)| respectively.

### System setup and simulation protocol

The sequence of the 672-bp DNA minicircle employed in this work has been based upon the template reported in Ref. [42] (see Section 1 of the Supplementary Material). To assess the role of the diverse topological regimes on the kinetics of the DNA linearization process, three levels of (initial) superhelical density have been characterized, namely *σ*_0_ = −0.06, 0, 0.06: These values belong to an average biological scenario [15], and are associated with a low likelihood of DNA denaturation defects and bubbles, as discussed elsewhere [43–45].

MD replicates have been carried out *via* the single GPU-power oxDNA simulation code [46, 47], and performed in the NVT ensemble, at a temperature T = 310 K and monovalent salt concentration of 0.15 M. A tailored simulation protocol has been adopted to estimate the probability distribution of the DNA linearization: As suggested in Ref. [33], this process is described by an exponential probability density function:

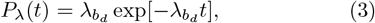

with 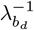 the characteristic time of the rupturing of DNA by a DSB motif at distance *b*_*d*_. As such, multiple trajectories are generally required to aptly describe the distribution of linearization events—and the statistical moments thereof. Within this protocol, each independent MD replicate has thus been set up according to a stepwise procedure as follows:

1. A perfectly circular, starting configurations of the DNA minicircle is generated *via* the tacoxDNA software [48] at a given *σ*_0_, by homogeneously distributing the excess twist (or the lack thereof) along the DNA filament. This structure is subjected to a fast MD relaxation stage of 1 × 10^6^ MD steps at T = 310 K employing the Bussi thermostat [49]: A modified version of the DNA backbone potential is enabled, offering a broader stability against the larger displacements of the particles. As suggested by the oxDNA developers, the correlation time and timestep frequency of the thermostat have been set to 1000 and 53 MD steps respectively, and associated with a simulation timestep Δ*t* = 9.09 fs (details of the conversion factor between oxDNA and SI units are reported in the work of Sengar and co-workers [32]);
2. A subsequent equilibration run of 3 × 10^7^ MD steps is thus performed employing the Langevin thermostat—the diffusion coefficient and timestep frequency set to 2.5 and 103 respectively, according to the indications of the oxDNA developers. The overall time of the MD simulation has been established upon (the decay of) the autocorrelation of the gyration radius of the DNA minicircle at the diverse degrees of superhelical density (reported in Ref. [43]);
3. At this stage, one of four DSB motifs (defined by the distance *b*_*d*_ = 0, 1, 2, 3 bps) is enforced on either an apical loop or the supercoiled region of the minicircle (as shown in Figure 1). To this purpose, an in-house script has been developed to generate the modified topology and configuration files of the broken DNA molecule in the oxDNA format. We note that DNA molecules with superhelical density value *σ*_0_ = 0 do not exhibit marked looped/supercoiled domains, on average: As such, DSB lesions have been placed at random on the DNA structure according to a uniform probability distribution for this sole scenario;
4. As the lesion is set, a MD simulation is performed employing the Langevin thermostat at T = 310 K. The sampling time of the individual replicate (*t*_*i*_) is defined within a discrete set of values homogeneously distributed within the range [0, T (*b*_*d*_)], with 𝒯 (*b*_*d*_) the timescale in which a DNA linearization event occurs with a 99% chance—according to Equation 3 and the values of 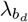 reported in Ref. [33];

- At the end of the MD replicates, the residual, non-bonded energy contributions between all nucleotides involved with the lesion site (i.e., located between and about the two backbone nicks—see Figure 1) are employed as proxy of the structural integrity of the DNA molecule. These are defined by:
- 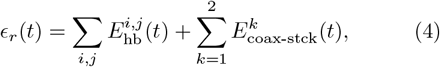

where 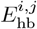 is the pairwise hydrogen bonding interaction between nucleotides, and 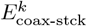 is the coaxial stacking energy between the nucleotides at the opposite sides of each nick. A DNA minicircle is irreversibly ruptured at the end of a MD trajectory if *ϵ*_*r*_(*t*) = 0.

We carried out 50 independent MD replicates for each combination of DSB motif, MD simulation time *t*_*i*_, and DNA lesion site (i.e., loop/supercoiled domain). Thus, the probability of DNA linearization at time *t*_*i*_ for a molecule characterized by an initial value of *σ*_0_—denoted *p*_lin_(*t*; *σ*_0_) throughout the manuscript—has been estimated for all DSB motifs and superhelical density regimes as the fraction of simulations scoring a rupturing event. Lastly, we shall remark that all time-related observables have been converted from oxDNA to SI units according to the rescaling factor reported in Ref. [32].

## RESULTS AND DISCUSSION

**Topological and conformational effects on the kinetics of the DNA linearization**

In a recent *in silico* characterization of the behavior of a linear DNA molecule, we have inferred that the kinetics of the DNA rupturing by DSBs follows an Arrhenius-like increase as function of the DSB distance *b*_*d*_ [33]. As such, we firstly focused on the statistics underlying the linearization process of the DNA minicircles subjected to diverse combinations of DSB motifs, lesion sites, and superhelical density.

We hereby define *p*_lin_(*t*; *b*_*d*_, *σ*_0_) as the probability to observe a rupturing event induced by a DSB motif of distance *b*_*d*_, in a DNA molecule characterized by an initial superhelical density *σ*_0_: Since the process is Poissonian [33], we performed a fitting procedure on the time evolution of *p*_lin_(*t*; *b*_*d*_, *σ*_0_) *via* an exponential probability distribution of the form:

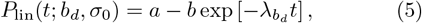

with *a, b* and 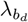 fitting parameters. Particularly, 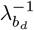 is the characteristic rupturing time associated with a DSB motif of distance *b*_*d*_.

Figure 2 shows the probabilities *P*_lin_(*t*; *b*_*d*_, *σ*_0_) associated with the diverse combinations of DSB motif and site, and initial superhelical density—while the outcome of the fitting procedure is reported in Table I. These data corroborate our earlier findings and highlight how topological features overall modulate the characteristic rupturing times (i.e.,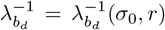). In fact, the mechanical strain enforced by the (excess/defect) DNA supercoiling enhances the linearization process but in the simulation scenario at [*σ*_0_ = −0.06, *r* = supercoiled], whose values of 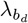 are close to a torsionally-relaxed regime (i.e., *σ*_0_ = 0).

**FIG. 2.**
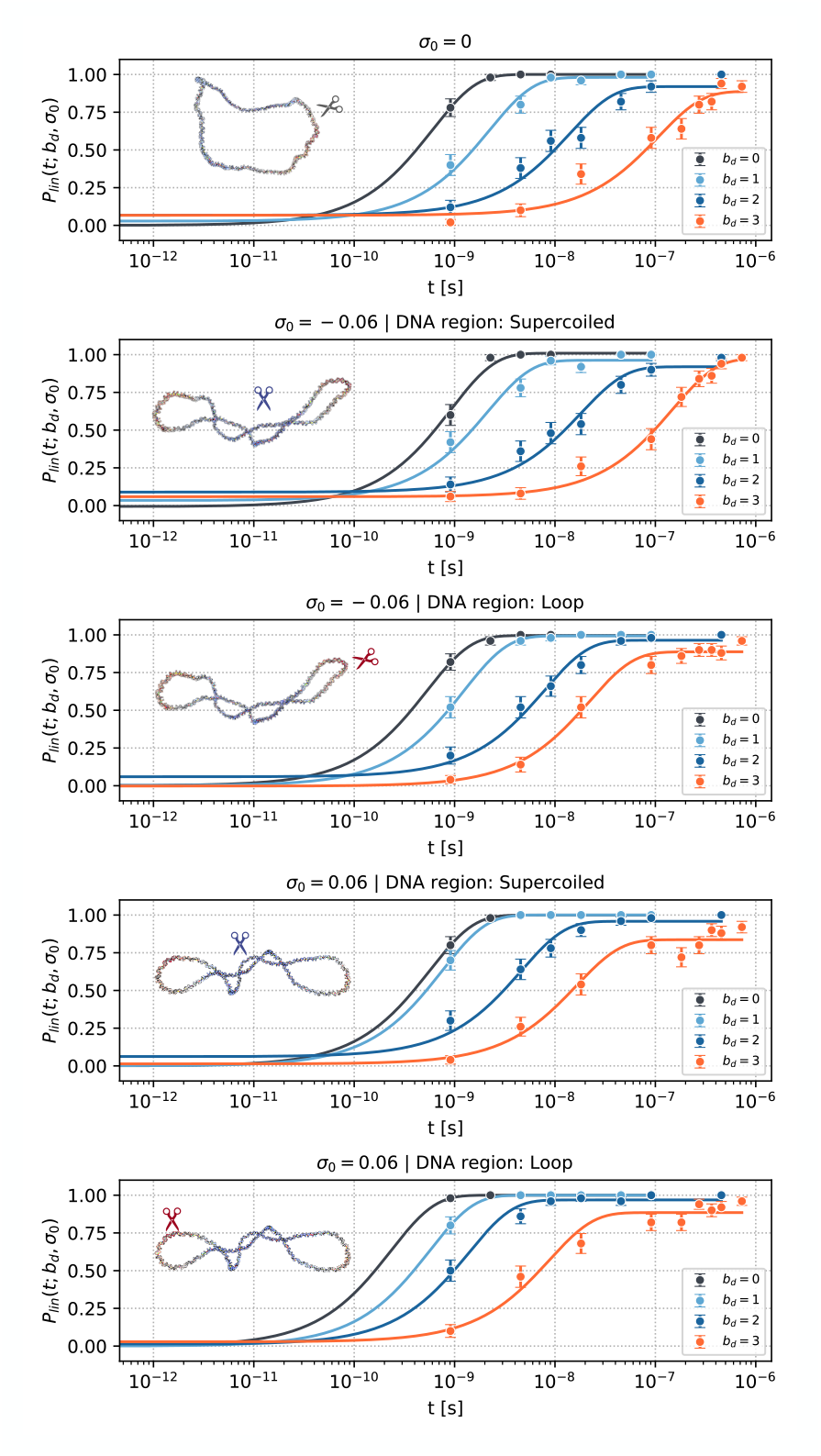
Outcome of the fitting procedure (based on Equation 5) of the rupturing probabilities of the 672-bps DNA minicircle, at each value of *t*_*i*_ and for diverse combinations of DSB motif (i.e., *b*_*d*_ = 0, 1, 2, 3), lesion site, and initial superhelical density *σ*_0_. The uncertainty associated with each point is defined upon the standard deviation of the binomial distribution, taking into account fifty independent MD replicates per scenario.

**TABLE I.** Parameters of the fitting procedure (based on Equation 5) of the DNA rupturing probabilities, for diverse combinations of DSB motif (i.e., b_d_ = 0, 1, 2, 3), lesion site, and initial superhelical density σ_0_ of the DNA minicircle.

| $b_d$ | $\lambda_{b_d}^{-1}(\sigma, r)$ [s] | | | | |
| --- | --- | --- | --- | --- | --- |
| | $\sigma_0 = 0$ | $\sigma = -0.06$ | | $\sigma = 0.06$ | |
|  | - | Supercoiled | Loop | Supercoiled | Loop |
| 0 | $(6.00 \pm 0.02) \times 10^{-10}$ | $(9.2 \pm 0.9) \times 10^{-10}$ | $(5.3 \pm 0.2) \times 10^{-10}$ | $(5.65 \pm 0.02) \times 10^{-10}$ | $(2.3239 \pm 0.0008) \times 10^{-10}$ |
| 1 | $(2.2 \pm 0.3) \times 10^{-9}$ | $(2.1 \pm 0.5) \times 10^{-9}$ | $(1.24 \pm 0.04) \times 10^{-9}$ | $(7.56 \pm 0.03) \times 10^{-10}$ | $(5.649 \pm 0.003) \times 10^{-10}$ |
| 2 | $(1.4 \pm 0.4) \times 10^{-8}$ | $(1.8 \pm 0.5) \times 10^{-8}$ | $(8 \pm 1) \times 10^{-9}$ | $(4.6 \pm 0.8) \times 10^{-9}$ | $(1.4 \pm 0.2) \times 10^{-9}$ |
| 3 | $(1.1 \pm 0.3) \times 10^{-7}$ | $(1.5 \pm 0.2) \times 10^{-7}$ | $(2.2 \pm 0.3) \times 10^{-8}$ | $(1.6 \pm 0.4) \times 10^{-8}$ | $(9.0 \pm 2.0) \times 10^{-9}$ |

To quantify the kinetic effect induced by the degree of supercoiling and the DSB site, we define the modulation factor of the linearization probability *K*(*t, σ, r*) as:

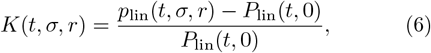

where the values *P*_lin_(*t*, 0) are obtained *via* Equation 5, with parameters derived from the fitting procedure. According to this definition, *K*(*t, σ, r*) offers an estimate of the kinetic enhancement across the significant timescales 𝒯 (*b*_*d*_) of the DNA linearization process, for each DSB scenario. In fact, as illustrated by Figure 3, the choice of the lesion site, as well as of the superhelical density regime, entails a time-dependent modulating effect in the kinetics of the DNA rupturing process, with the positive supercoiling inducing a stronger enhancement, irrespective of the DNA lesion site. Additionally, within each initial superhelical density regime, DSBs placed on apical loops are more likely to induce the failure of the residual nonbonded interactions leading to the rupturing of the DNA molecule.

**FIG. 3.**
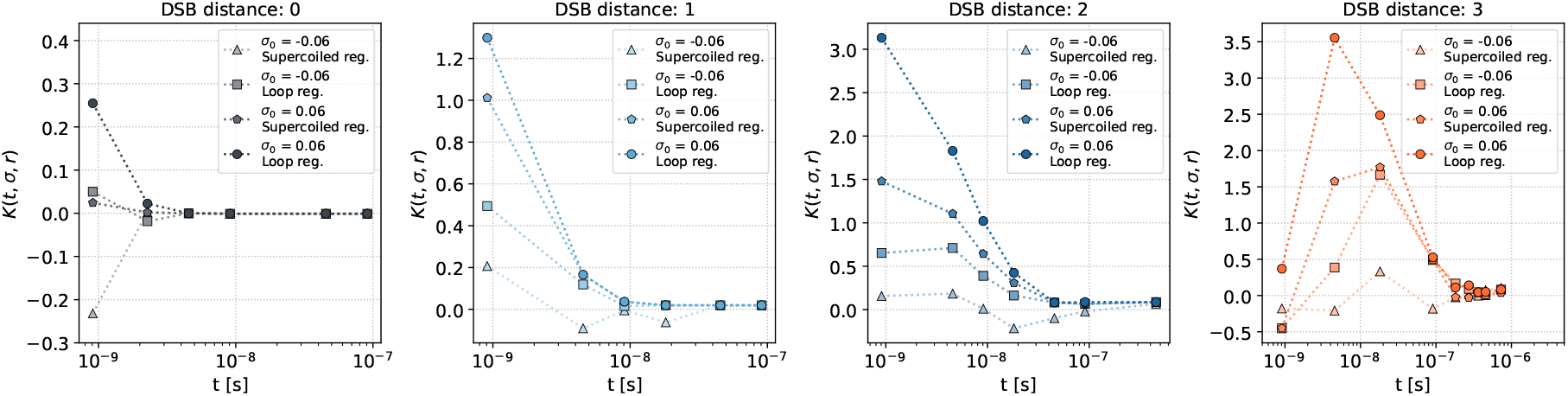
Modulation factor of the rupturing probability (with respect to a torsionally-relaxed scenario, i.e., *σ*_0_ = 0), for each combination of superhelical density and lesion site (details in the text).

We shall note that if the kinetic effect associated with the localization of the DNA lesion is somewhat intuitive—with the sharply-bent apical loops enforcing a significant structural distortion that favors the linearization process—the outcome associated with the diverse degrees of DNA supercoiling is less trivial. Positively supercoiled DNA molecules show a higher likelihood of structural failure: This might be accounted for by the lack of an effective release pathway of the excess torsional stress in the free energy landscape. Conversely, the partial unwinding of the DNA molecule under a negative supercoiling regime causes a mild enhancement of the rupturing probability by DSBs located at apical loops, and no detectable enhancement in the supercoiled domain—if any, a stabilizing effect on the DNA structure is appreciated. In fact, despite both positivelyand negativelysupercoiled regimes entailing a similar level of structural compaction of the DNA molecule—which is seemingly associated with a reduced susceptibility of DNA towards certain radiation qualities [22]—the latter does not quite affect the kinetics of the rupturing process, and specifically so in the supercoiled domain of a closed DNA moiety.

### Time evolution of the topological features in lesioned DNA

By disrupting the backbone connectivity, (single, double) DNA strand breaks compromise the topological invariance of the superhelical density, triggering the torsional relaxation of the DNA molecule. In the light of the time-dependency of the modulation factor *K*, we thus characterized the time evolution of the superhelical density from an initial value *σ*_0_, on DNA minicircles broken by DSB motifs that are sufficiently stable (that is, at distance *b*_*d*_ = 2, 3). To this aim, we carried out several, finely-sampled, MD replicates, following a similar step-wise protocol as that described in the Materials and Methods section, for an overall simulation time of about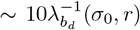: The behavior of *σ*(*t*) has been evaluated on stable DNA conformations, (i.e., preserving all stacking and hydrogen-bonding interactions of the native pairing conformations), by computing the DNA twist and writhe according to Klenin and Langowski [50]. Figure S1 and Figure S2 shows the evolution of *σ* for the DSB scenarios at distance 2 and 3: The disruption event triggers a mechanical relaxation, whereby the DNA releases the (excess) torsional stress *via* an over/under-winding process in the negative and positive superhelical density regimes respectively, in line with earlier works [51, 52]. Additionally, Figure 4 shows the time evolution of the average linking difference ⟨ΔLk⟩ = ⟨Lk(*t*) − Lk_0_⟩ (taking into account all independent MD replicates)—defining the average number of helical turns released (or rewound) by the minicircle—with Lk_0_ the linking number of a topologically-relaxed scenario. We observe that the over/under-winding process lowers ⟨ΔLk⟩ by about ±0.5 turns within a timescale defined by 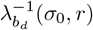 (corresponding to a DNA linearization probability of 63%). Thus, despite the mechanical relaxation, the initial superhelical density values are preserved on average. Arguably, the relaxation process itself might contribute to the structural destabilization of the DNA molecule, with no MD replicates reaching supercoiling values of Lk_0_ beyond the characteristic rupturing timescale (see Figure S3 in the Supplementary Material). Overall, these results are well summarized by the behavior of *K*(*t, σ, r*) whose value is ultimately defined by the multiple, concurrent processes involved with the DNA rupturing. Particularly, within a short timescale (i.e., up to about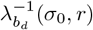), the DNA rupturing probability is seemingly insensitive to large-scale structural changes —as suggested by Figure 4 alike. Therefore, *p*_lin_ is mostly determined by local structural features of the molecule, such as the sequence of nucleotides involved with the DSB, the (initial) levels of mechanical stress, and the location of the lesion. Arguably, *K* should converge to a constant value within this time range, given by:

**FIG. 4.**
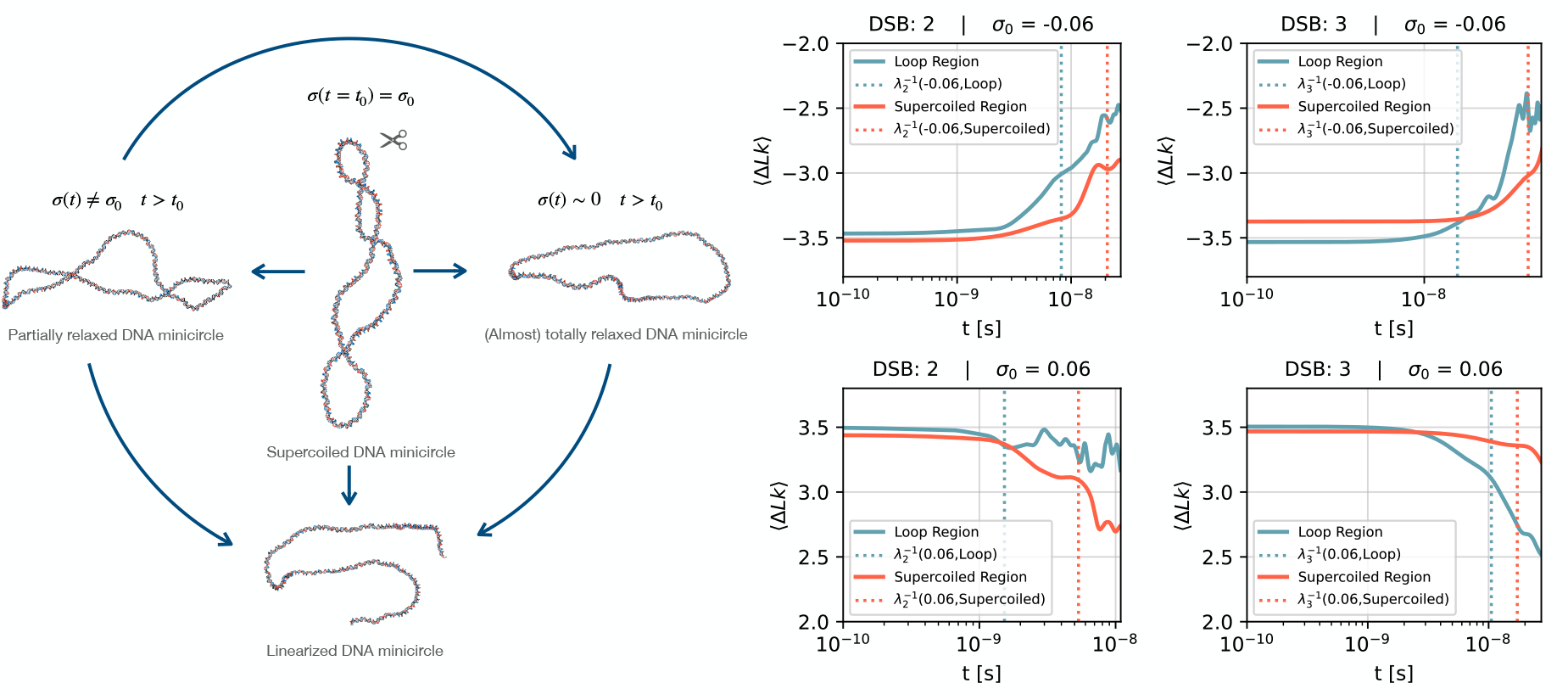
(**Left**) Schematic depiction of the processes unfolding throughout the evolution of a broken DNA minicircle—namely, the relaxation of the (excess) torsional stress and the linearization of the molecule. (**Right**) Time evolution of the linking difference, averaged over the independent MD replicates, at initial superhelical density *σ*_0_ = *±*0.06. Data refer to the DSB scenarios at distance 2 and 3, enforced on diverse lesion sites (i.e., either apical loops or the supercoiled region). The vertical lines mark the characteristic DNA rupturing times 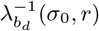 associated with each scenario.

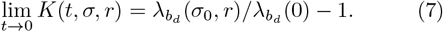

On an intermediate timescale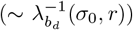, the structural dynamics of DNA is affected by the lesion itself, which triggers the release of torsional stress *via* an over/under-winding process. Throughout this process, the DNA molecule eventually converges to a torsionally-relaxed conformation identified by *σ* ∼ 0, if the residual interactions holding the DNA molecule about the lesion site are sufficiently stable.

Lastly, in the long-time regime, the linearization probability approaches *P*_lin_(*t*, 0), irrespective of the (initial) superhelical density regime.

### Topological and structural effects on the reaction pathway of the DNA linearization process

So far, we have discussed how the initial, topological state, as well as the DSB lesion site, are key variables modulating the kinetics of the DNA linearization process. This observation is recapitulated by the analysis of the free energy profile of the DNA rupturing by DSB motifs at distances of 2 and 3, defined by tracking the evolution of the residual interactions about the lesion site over time.

Starting from lesioned, equilibrated DNA conformations, we thus performed several finely-sampled MD replicates for each supercoiling scenario, lesion site and DSB motif, for an overall simulation time of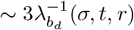.

Figure 5 and Figure S4 illustrate the free energy landscapes (expressed in internal oxDNA energy units) of the DNA rupturing process for the DSB scenarios at distance *b*_*d*_ = 3: As putative reaction coordinates describing the DNA rupturing process, we employed i) the sum of the residual coaxial stacking interactions between the nucleotides at both DNA termini (Ψ_*stck*_), and ii) the sum of the hydrogen-bonding interactions between all nucleotide pairings within the twofold nick of the DNA backbone (Ψ_*HB*_). As similar observations apply to DSB scenarios at distance *b*_*d*_ = 2 and 3 (the free energy landscapes for the former are shown in Figure S5), we will hereby focus the discussion on the latter, for the sake of clarity.

**FIG. 5.**
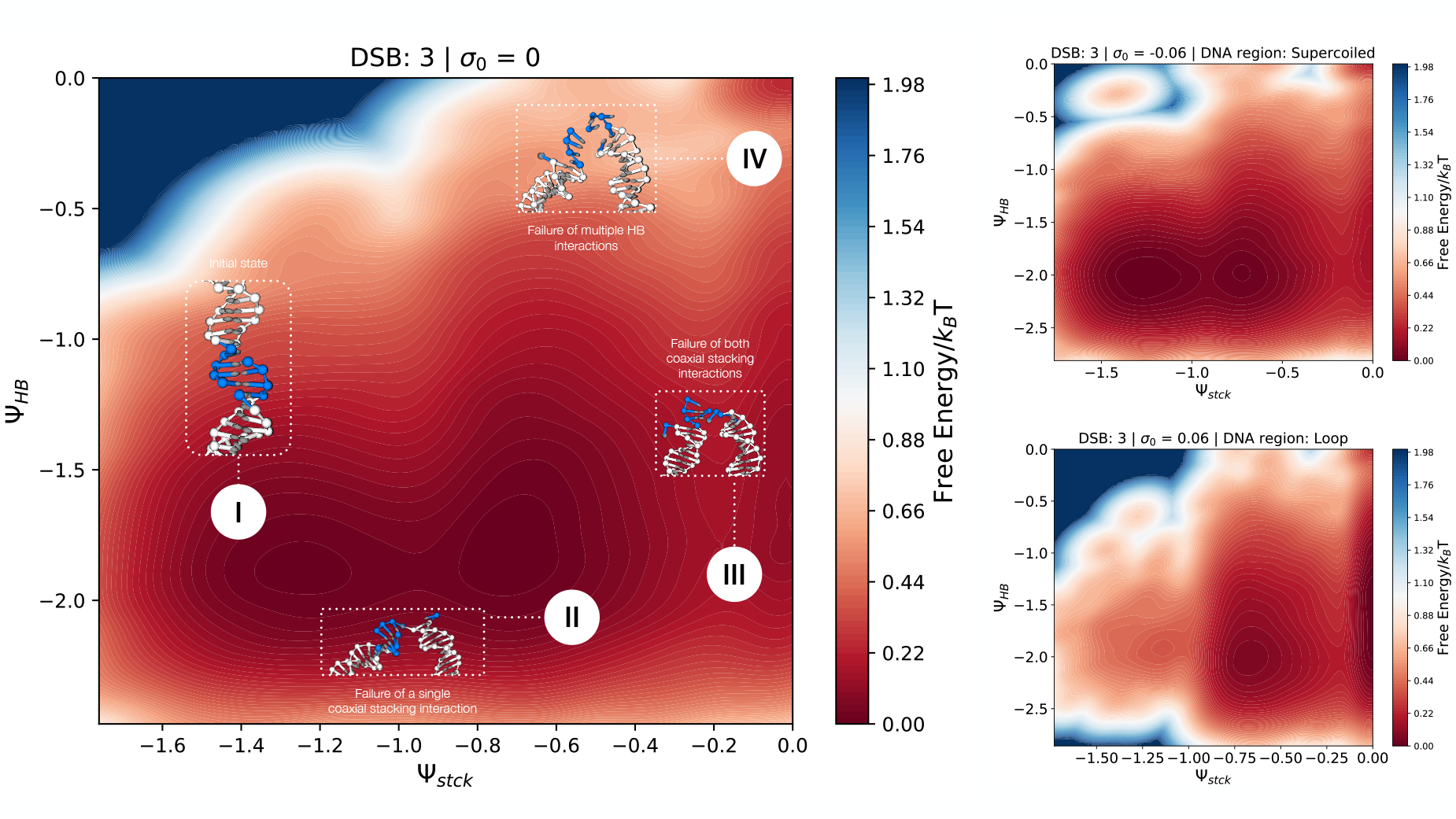
Free energy landscapes depicting the rupturing process of a DNA minicircle broken by a DSB motif at distance 3, at diverse combinations of (initial) superhelical density and lesion site. The putative reaction coordinates correspond to the sum of the hydrogen-bonding interactions of all nucleotide pairings in-between the twofold DNA nick (Ψ_*HB*_), and the sum of the coaxial stacking interactions between nucleotides at the broken DNA termini (Ψ_*stck*_)—both coordinates are expressed in oxDNA internal units. Four thermodynamic basins have been identified, numbered **I** through **IV**, associated with the progressive loosening of the non-covalent bonds at the lesion site (details in the main text).

Each free energy landscape is characterized by four major thermodynamic basins:

- The first basin (**I**) is located in the lower-left region of the free energy landscape, and identifies a thermodynamic state of the DNA molecule where all coaxial stacking and hydrogen-bonding interactions are intact;
- The second basin (**II**) is located in the mid-lower region, and identifies a metastable state of the DNA molecule where a single coaxial stacking interaction is broken, while promoting the exploration of the free energy landscape along Ψ_*HB*_. Arguably, this basin is populated by DNA configurations emerging during the over/under-winding process;
- The third basin (**III**) lies in the lower-right region of the landscape: Here, the residual coaxial stacking energy is zero, and the lesion site is held solely by (highly-fluctuating) hydrogen-bonding interactions;
- Lastly, the fourth basin (**IV**) is located in the upper-right corner, identifying a thermodynamic state where no residual interactions hold the DNA termini, driving the system towards a fully-linearized configuration.

By benchmarking all free energy landscapes against the scenario at *σ*_0_ = 0, we observe that the excess mechanical stress stored in the DNA molecule modulates the shape of the four thermodynamics basins—and the relative populations thereof.

Basin **II** often broadens along the Ψ_*HB*_ reaction coordinate, showing a higher population with respect to basin **I**. Scenarios [*σ* = 0.06, *r* = supercoiled] and [*σ* = −0.06, *r* = loop] exhibit similar profiles, with basins **II** and **III** broadly merging in the former case—in line with the enhanced rupturing effectiveness, quantified by the *K* factor.

Conversely, scenarios [*σ* = −0.06, *r* = supercoiled] and [*σ* = 0.06, *r* = loop] show completely different free energy landscapes. Basin **I** is highly populated in the former, highlighting a significant structural stability: Moreover, this scenario features a small basin in the upper-left corner of the landscape, possibly identifying a denatured DNA state of sorts, which is observed in the [*σ* = −0.06, *r* = loop] scenario alike. In the [*σ* = 0.06, *r* = loop] scenario, basin **I** (defining the structural stability of the lesion site) is nearly absent, whereas **II** and **III** are the most populated basins. This instability agrees with the results illustrated in Figure 3: While this behavior is less pronounced in the DSB scenario at *b*_*d*_ = 2 (see Figure S5 of the Supplementary Material), a population imbalance between basins **I** and **II** is consistently observed under positive superhelical density regimes.

According to this qualitative depiction, the DNA rupturing pathway is seemingly triggered by the disruption of (coaxial) stacking interactions: Possibly, the failure of a single nicking site underlies a multitude of destabilizing pathways, i.e., triggering the torsional relaxation of the DNA molecule and/or enhancing the localized fraying of the DNA termini at the lesion interface. Subsequent to the failure of the second nicking site, the rupture of the hydrogen bonds unfolds *via* a highly-cooperative process, seemingly following an unzipping mechanism, as observed in Ref. [33].

## CONCLUSIONS

Double strand breaks (DSBs) involve the rupture of the covalent backbone of the DNA helix on both complementary strands, and are a detrimental outcome of the action of external vectors—such as radicals and ionizing radiations—as well as of a variety of cellular routines. Earlier works have implied that diverse DSB motifs (i.e., associated with a variable distance between the twofold nick of the DNA backbone, *b*_*d*_) bear different kinetic and thermodynamic implications, potentially impacting on the structural dynamics of the DNA molecule [28, 33].

In this work, we have assessed the interplay between structural and topological features of the DNA molecule—namely, the location of the lesion site and the (initial) level of superhelical density—in modulating the kinetics of the linearization of a 672-bp DNA minicircle, subjected to diverse DSB motifs at *b*_*d*_ = 0, 1, 2, 3, by means of classical, coarse-grained, molecular dynamics simulations.

Firstly, we observed that DSB lesions exhibit a significantly higher rupturing probability on apical loops than on highly-writhed supercoiled regions—the former being more inclined to mechanical failures, yet associated with a lower effective target volume [21, 22].

We thus examined the influence of positive and negative supercoiling on the characteristic times of the DNA rupturing process, as opposed to a topologically-relaxed, reference system—i.e., at zero superhelical density. While both regimes overall enhance the probability of DNA linearization, the effect is significantly stronger under positive supercoiling. This asymmetry is recapitulated by the free-energy landscapes of the rupturing process, described upon the evolution of the (coaxial-stacking, hydrogen-bonding) residual interactions between the nucleotides involved with the lesion site: In fact, these profiles show that positive supercoiling induces a stronger destabilization of the DNA helix, favoring configurations that feature the failure of at least one nicking site.

The simulation of sufficiently stable DSB motifs reveals the existence of a complex, time-dependent interplay between structural and mechanical features of the DNA molecule, on account of *σ* no longer being a topological invariant, subsequent to the (twofold) nicking of the DNA backbone. Indeed, the DNA minicircle rewinds/unwinds the excess torsional stress, eventually converging to a relaxed mechanical state. However, this process arguably destabilizes the structural integrity of the DNA helix, contributing to the molecular rupture.

The results discussed in this work offer valuable insights in the context of DNA radiosensitivity. Notably, simulations of DSB lesions enforced over the supercoiled region of a minicircle at *σ*_0_ = −0.06 (which is a value of biological significance [15]) show no kinetic enhancement in the linearization probability of the DNA molecule—while exhibiting a free energy landscape that is close to a reference scenario at *σ*_0_ = 0. In fact, we note that, despite the topologically-relaxed minicircle exhibiting the slowest characteristic times of the DNA rupturing, it might not be the least radiosensitive. As discussed in earlier works alike [6, 7, 21, 22], compact DNA conformations (as those achieved by supercoiled DNA molecules) contribute to its radiation resistance by lowering the effective volume of the target DNA. In line with these observations, we corroborate the idea that biologically-significant levels of negative superhelical density (e.g., modulating the expression, regulation, and compartmentalization of the genome) might bear a further level of radiation protection at a molecular scale, from both a kinetic and a structural viewpoint.

## AUTHOR CONTRIBUTIONS

LP and RP oversaw and coordinated the whole research project. MM performed all MD simulations. MM and LP designed the analysis protocol: The data analysis was carried out by MM with the constant supervision of LP. MM and LP mainly worked on the first draft, while the final manuscript was written with contributions from all authors.

## SUPPLEMENTARY MATERIAL

Supplementary Material is available. Moreover, the raw data and scripts supporting the findings of this work are available from the corresponding author upon reasonable request.

## ACKNOWLEDGMENTS

The authors acknowledge support from the following Agencies: CINECA award under the ISCRA initiative, for the availability of high-performance computing resources; Fondazione CARITRO through the project COMMODORE (#20260); the Italian Ministry of Education, University and Research (MIUR) through the FARE grant for the project HAMMOCK (grant R18ZHWY3NC); ICSC – Centro Nazionale di Ricerca in HPC, Big Data and Quantum Computing, funded by the European Union under NextGenerationEU; PRIN 2022 PNRR Prot. n. 2022Z3BBPE. Views and opinions expressed are however those of the author(s) only and do not necessarily reflect those of the European Union or The European Research Executive Agency. Neither the European Union nor the granting authority can be held responsible for them.

