## Supplementary Material for "Dynamical insights on the role of supercoiling on DNA radiosensitivity"

#### 1 DNA template

Here is reported the 5'-3' sequence of the 672-bps DNA minicircle employed in this work, based on the template described by Fogg and co-workers [1].

```
  1 TTTATACTAA CTTGAGCGAA ACGGGAAGGG TTTTCACCGA TATCACCGAA
 51 ACGCGCGAGG CAGCTGTATG GCATGAAAGA GTTCTTCCCG GAAAACGCGG
101 TGGAATATTT CGTTTCCTAC TACGACTACT ATCAGCCGGA AGCCTATGTA
151 CCGAGTTCCG ACACTTTCAT TGAGAAAGAT GCCTCAGCTC TGTTACAGGT
201 CACTAATACC ATCTAAGTAG TTGATTCATA GTGACTGCAT ATGTTGTGTT
251 TTACAGTATT ATGTAGTCTG TTTTTTATGC AAAATCTAAT TTAATATATT
301 GATATTTATA TCATTTTACG TTTCTCGTTC AGCTTTTTTA TACTAACTTG
351 AGCGAAACGG GAAGGGTTTT CACCGATATC ACCGAAACGC GCGAGGCAGC
401 TGTATGGCAT GAAAGAGTTC TTCCCGGAAA ACGCGGTGGA ATATTTCGTT
451 TCCTACTACG ACTACTATCA GCCGGAAGCC TATGTACCGA GTTCCGACAC
501 TTTCATTGAG AAAGATGCCT CAGCTCTGTT ACAGGTCACT AATACCATCT
551 AAGTAGTTGA TTCATAGTGA CTGCATATGT TGTGTTTTAC AGTATTATGT
601 AGTCTGTTTT TTATGCAAAA TCTAATTTAA TATATTGATA TTTATATCAT
651 TTTACGTTTC TCGTTCAGCT TT
```

### 2 Parameters of the fitting of the linearization probabilities

| | | | | | |
|---|---|---|---|---|---|
| | | $\sigma_0 = 0$ | | | |
| $b_d$ | $a$ | $b$ | $\lambda_{b_d}^{-1}$ [s] | Adj. $R^2$ | RMSE |
| 0 | $1.0005 \pm 0.0006$ | $1.0006 \pm 0.001$ | $(6.00 \pm 0.02) \times 10^{-10}$ | 1.00 | 0.001 |
| 1 | $0.98 \pm 0.02$ | $0.95 \pm 0.05$ | $(2.2 \pm 0.3) \times 10^{-9}$ | 0.98 | 0.03 |
| 2 | $0.92 \pm 0.06$ | $0.85 \pm 0.08$ | $(1.4 \pm 0.4) \times 10^{-8}$ | 0.93 | 0.07 |
| 3 | $0.89 \pm 0.05$ | $0.82 \pm 0.06$ | $(1.1 \pm 0.3) \times 10^{-7}$ | 0.94 | 0.07 |
| | | $\sigma = -0.06$ \| DNA region: Supercoiled | | | |
| $b_d$ | $a$ | $b$ | $\lambda_{b_d}^{-1}(\sigma, r)$ [s] | Adj. $R^2$ | RMSE |
| 0 | $1.01 \pm 0.01$ | $1.02 \pm 0.04$ | $(9.2 \pm 0.9) \times 10^{-10}$ | 0.99 | 0.02 |
| 1 | $0.96 \pm 0.03$ | $0.93 \pm 0.06$ | $(2.1 \pm 0.5) \times 10^{-9}$ | 0.97 | 0.05 |
| 2 | $0.92 \pm 0.05$ | $0.83 \pm 0.07$ | $(1.8 \pm 0.5) \times 10^{-8}$ | 0.94 | 0.06 |
| 3 | $0.97 \pm 0.04$ | $0.92 \pm 0.04$ | $(1.5 \pm 0.2) \times 10^{-7}$ | 0.98 | 0.04 |
| | | $\sigma = -0.06$ \| DNA region: Loop | | | |
| $b_d$ | $a$ | $b$ | $\lambda_{b_d}^{-1}(\sigma, r)$ [s] | Adj. $R^2$ | RMSE |
| 0 | $0.996 \pm 0.006$ | $1.00 \pm 0.01$ | $(5.3 \pm 0.2) \times 10^{-10}$ | 1.00 | 0.009 |
| 1 | $0.993 \pm 0.004$ | $0.99 \pm 0.01$ | $(1.24 \pm 0.04) \times 10^{-9}$ | 1.00 | 0.007 |
| 2 | $0.96 \pm 0.03$ | $0.91 \pm 0.05$ | $(8 \pm 1 \times 10^{-9}$ | 0.97 | 0.05 |
| 3 | $0.89 \pm 0.02$ | $0.89 \pm 0.03$ | $(2.2 \pm 0.3) \times 10^{-8}$ | 0.98 | 0.04 |
| | | $\sigma = 0.06$ \| DNA region: Supercoiled | | | |
| $b_d$ | $a$ | $b$ | $\lambda_{b_d}^{-1}(\sigma, r)$ [s] | Adj. $R^2$ | RMSE |
| 0 | $0.9997 \pm 0.0005$ | $1.000 \pm 0.001$ | $(5.65 \pm 0.02) \times 10^{-10}$ | 1.00 | 0.001 |
| 1 | $1.0005 \pm 0.0005$ | $1.001 \pm 0.001$ | $(7.56 \pm 0.03) \times 10^{-10}$ | 1.00 | 0.001 |
| 2 | $0.96 \pm 0.03$ | $0.90 \pm 0.05$ | $(4.6 \pm 0.8) \times 10^{-9}$ | 0.97 | 0.05 |
| 3 | $0.84 \pm 0.03$ | $0.82 \pm 0.05$ | $(1.6 \pm 0.4) \times 10^{-8}$ | 0.96 | 0.06 |
| | | $\sigma = 0.06$ \| DNA region: Loop | | | |
| $b_d$ | $a$ | $b$ | $\lambda_{b_d}^{-1}(\sigma, r)$ [s] | Adj. $R^2$ | RMSE |
| 0 | $1.00001 \pm 0.00001$ | $1.00001 \pm 0.00003$ | $(2.3239 \pm 0.0008) \times 10^{-10}$ | 1.00 | 0.00002 |
| 1 | $1.00006 \pm 0.00006$ | $1.0006 \pm 0.0002$ | $(5.649 \pm 0.003) \times 10^{-10}$ | 1.00 | 0.0001 |
| 2 | $0.97 \pm 0.02$ | $0.96 \pm 0.04$ | $(1.4 \pm 0.2) \times 10^{-9}$ | 0.98 | 0.03 |
| 3 | $0.88 \pm 0.03$ | $0.86 \pm 0.06$ | $(9.0 \pm 2.0) \times 10^{-9}$ | 0.95 | 0.06 |

**Table S1:** Expanded version of Table 1 in the main text, showing the parameters of the fitting procedure of the DNA rupturing probabilities (based on Equation 4), for diverse combinations of DSB motif (i.e., $b_d = 0, 1, 2, 3$), lesion site, and initial superhelical density $\sigma_0$ of the DNA minicircle. The statistical estimators of the quality of the fitting procedure (i.e. the adjusted R-squared and the root mean square error) have been rounded up to the least significant digit.

### 3 Relaxation of the excess supercoiling in broken DNA minicircles

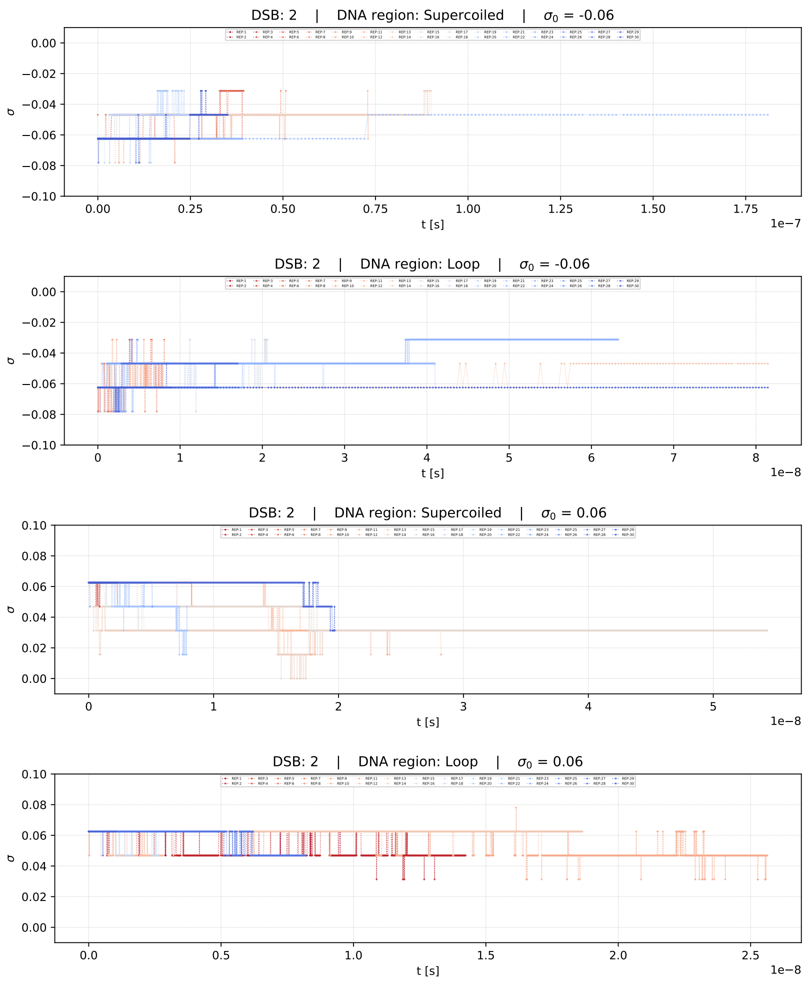


**Figure S1:** Time evolution, and relaxation, of the excess supercoiling levels in DNA minicircles broken by DSBs at distance $b_d = 2$, for diverse combinations of (initial) supercoiling regime and lesion site—each track is associated with a single MD replicate.

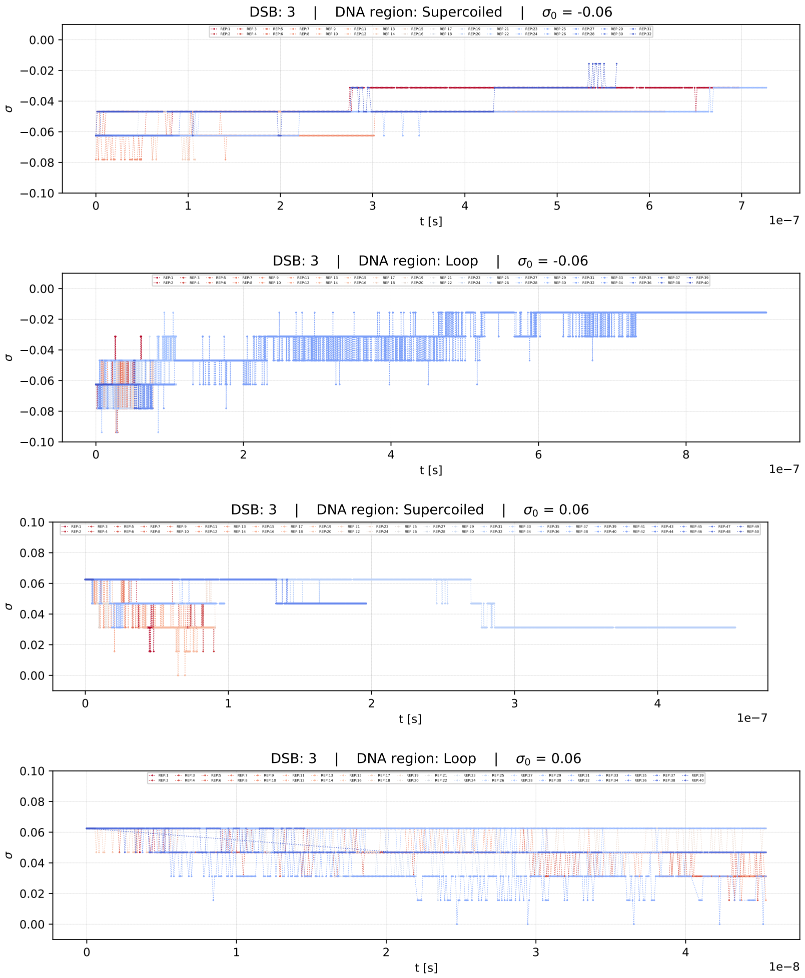


**Figure S2:** Time evolution, and relaxation, of the excess supercoiling levels in DNA minicircles broken by DSBs at distance $b_d = 3$, for diverse combinations of (initial) supercoiling regime and lesion site—each track is associated with a single MD replicate.

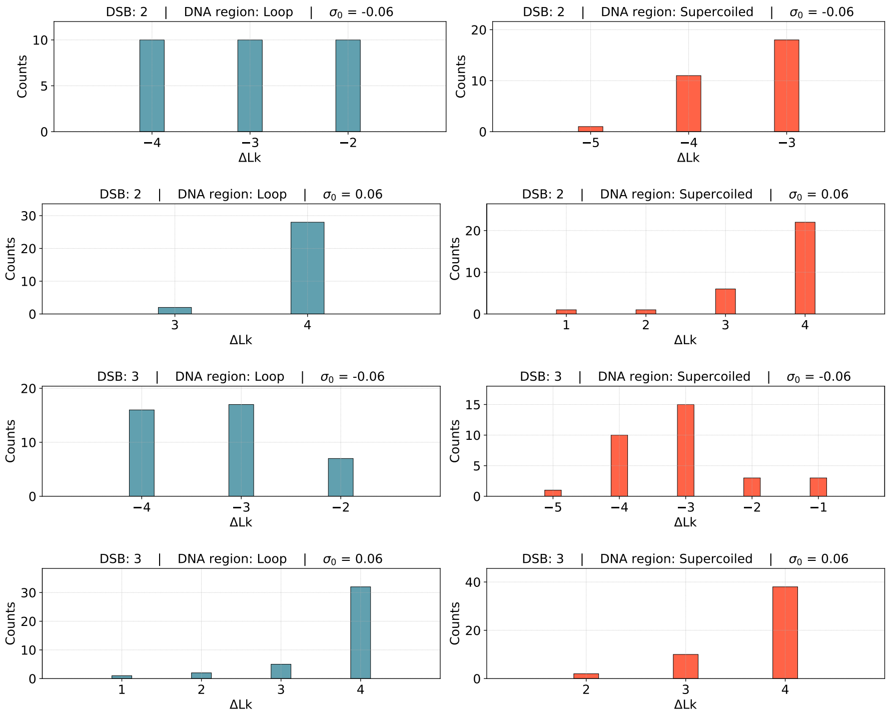


**Figure S3:** Distribution of the values of the linking differences ΔLk at the end of each independent MD replicate. The data refer to the DSB scenarios at distance 2 and 3, and diverse combinations of (initial) superhelical density and lesion site. The linking difference corresponding to a superhelical density of $|\sigma_0| = 0.06$ amounts to $|\Delta\text{Lk}| \sim 4$.

### 4 Free energy landscapes of the DNA rupturing

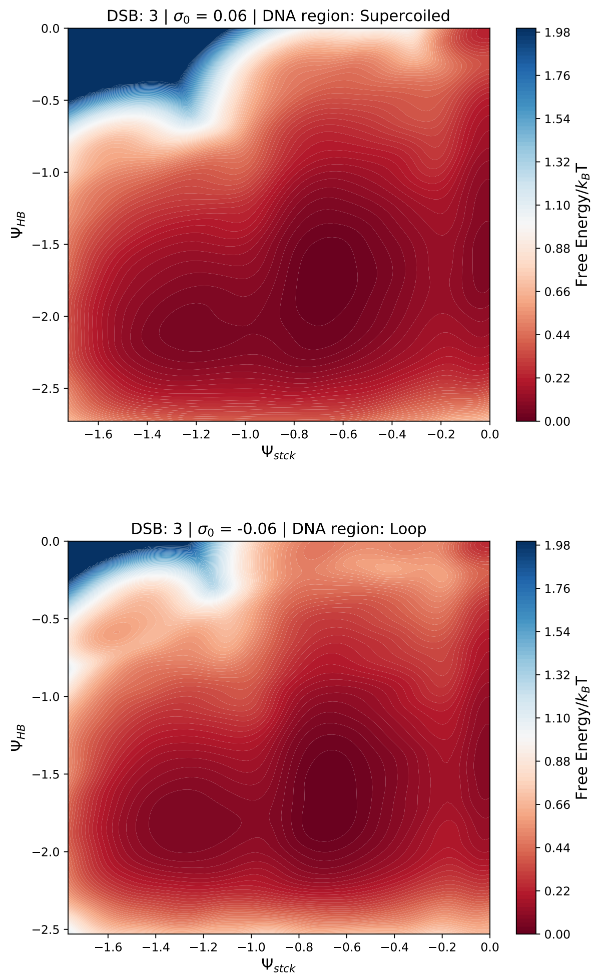


**Figure S4:** Free energy landscapes depicting the rupturing process of a DNA minicircle broken by a DSB motif at distance 3, associated with the [$\sigma_0 = 0.06$, *supercoiled*] and [$\sigma_0 = -0.06$, *loop*] scenarios (details in the main text).

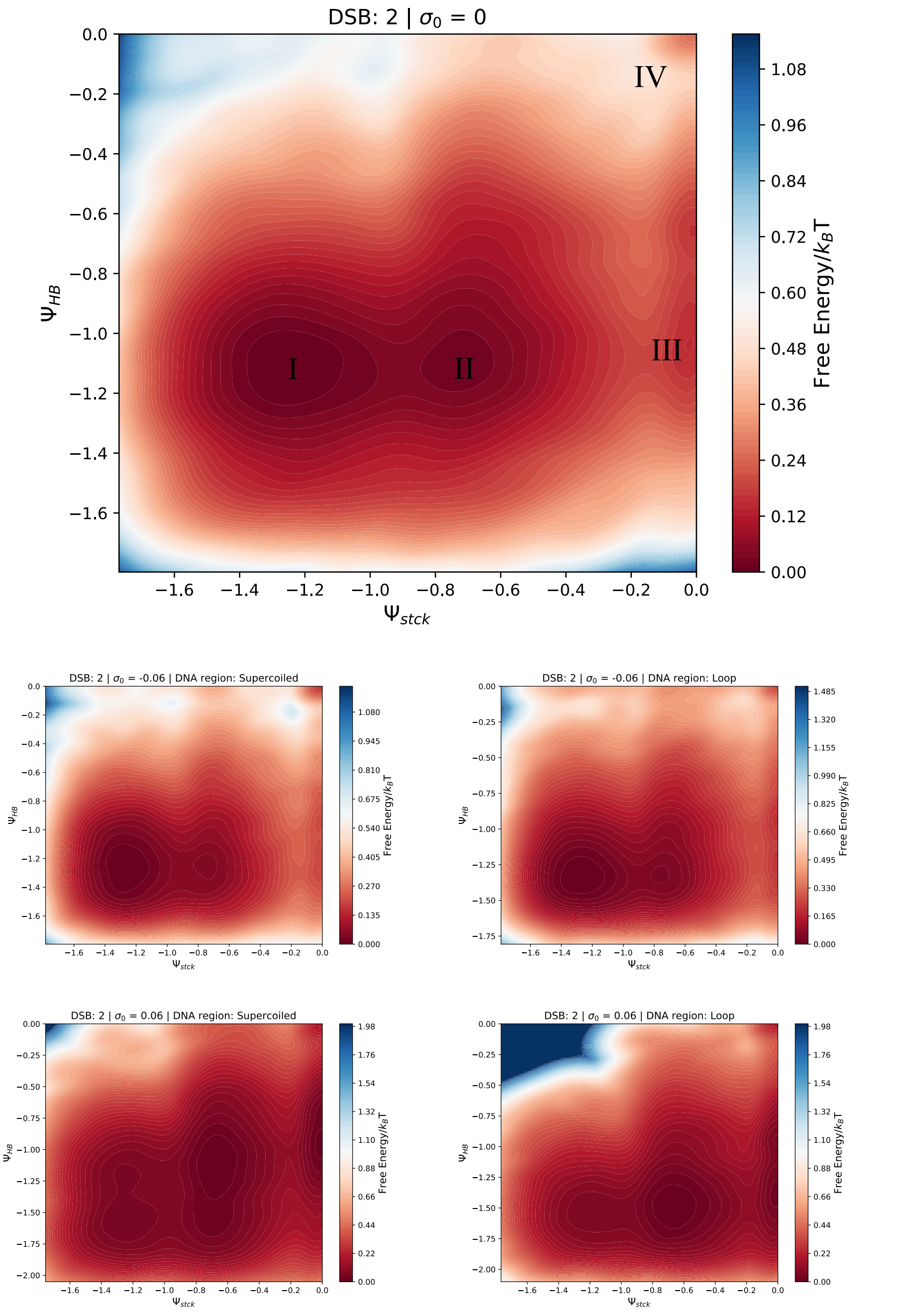


**Figure S5:** Free energy landscapes depicting the rupturing process of a DNA minicircle broken by a DSB motif at distance 2, for all combinations of (initial) supercoiling level and lesion site—details in the main text.
